# Microbial Food Safety in the Maryland Direct-to-Consumer Poultry Supply Chain

**DOI:** 10.1101/643106

**Authors:** Patrick A. Baron, David C. Love, Shanna Ludwig, Kathryn Dalton, Jesper Larsen, Gabriel K. Innes, Meghan F. Davis, Christopher D. Heaney

## Abstract

Direct-to-consumer food marketing is a growing niche in the United States food supply chain. Food animal producers who use direct marketing may employ different production models and standard practices from producers selling animal products to the conventional food system. Direct-to-consumer food supply chains (generally and specifically regarding food animal products) are relatively unexplored in food safety and health research. We conducted a cross-sectional, market-basket analysis of the Maryland direct-to-consumer poultry supply chain to assess food safety. We analyzed 40 direct-to-consumer commercial poultry meat products (one product per farm) for *Escherichia coli, Staphylococcus aureus* and *Salmonella spp.* using culture-based methods. Isolates underwent antimicrobial susceptibility testing. *E. coli* and *S. aureus* were recovered from 9/40 (23%) and 12/40 (30%) of poultry meat samples, respectively. Of interest for comparing direct-market and mainstream supply chains for food safety risks, no *Salmonella* isolates were recovered from any direct-market sampled poultry products and no multidrug resistance was observed in *E. coli* and *S. aureus* isolates. Microbial outcomes were compared to a survey of poultry production and processing practices within the same study population.

**Importance:** This study demonstrates substantially lower rates of antimicrobial-resistant (AMR) microbial pathogens in the market-basket products from Maryland direct-market broiler poultry supply chain compared to rates of AMR in the conventional supply chain for similar retail meat products from NARMS. We further describe the landscape of the statewide supply chain for direct-market poultry, focusing on characteristics related to risk management strategies applied to microbial food safety. These findings are of public health significance for both the research and policy communities; these data provide an initial evidence base for more targeted research evaluating potential risk factors for microbial food safety in the direct-to-consumer supply chain. These data will also assist the Maryland Department of Agriculture and other state-level agencies with oversight of food safety issues to guide policy efforts for direct-market poultry production and sales.

## 1. Introduction

*Escherichia coli, Salmonella spp.,* and *Staphylococcus aureus* are major causes of bacterial foodborne illness; however, US population exposure to these pathogens through non-industrial supply chains for livestock products is virtually unexplored in health and food safety research. The Centers for Disease Control and Prevention estimates that 1 in 6 people in the US acquire foodborne infections every year, with 128,000 hospitalizations and ~3,000 annual deaths [1]. Incidence of O157 and non-O157 Shiga-toxin producing *E. coli* (STEC) are estimated to cause illness at rates of 1.15 and 1.17 per 100,000, respectively [2]. *Salmonellosis* caused an estimated 1,027,561 cases of foodborne illness in 2013 in the US, resulting in ~19,000 hospitalizations and 380 deaths [3]. Other bacterial pathogens commonly associated with foodborne illness include *S. aureus* intoxication [3]. A review of food safety data from 1998-2008 indicates that poultry products contaminated with pathogenic bacteria comprised 17.9% of the annual burden of foodborne illness cases caused by bacterial exposure [4].

Industrial food animal production methods raise animals in high densities and producers often routinely use antimicrobials for disease prevention and therapeutic purposes [5–9], which may facilitate selection for antibiotic resistance among zoonotic bacteria. Antimicrobial resistance among foodborne bacterial pathogens is a complicating factor in foodborne illness; antimicrobial-resistant infections resulting from human exposure to foodborne bacteria caused an estimated 430,000 illnesses in the US in 2012 [10, 11]. The model(s) currently in use for direct-market poultry production have not been adequately investigated for their potential to facilitate selective pressure for antimicrobial resistance in foodborne pathogens.

The prevalence of microbial foodborne pathogens in consumer poultry meat products coming from the direct-market poultry supply chain remains relatively unexplored in health research. Some recent research has focused on the epidemiology of *Listeria* in the production environments of direct-to-consumer farms [12] and of *Salmonella spp.* in pastured-poultry production [13]. Only a handful of studies have evaluated microbial food safety risks in direct-market poultry supply chains [14, 15]; only one study addressed these issues through a market-basket and consumer exposure research lens [16]. This single study contained several methodological limitations which limit the interpretation of these findings (see Supplement).

The current study addresses the research gaps surrounding microbial food safety of direct-marketing systems for poultry in Maryland and builds on qualitative research in this population which demonstrated that the models, practices and inputs used in Maryland direct-market poultry production depart substantially from the typical models and practices of industrial-scale poultry production [17]. We therefore hypothesized that these inter-supply chain differences contribute to different microbial food safety outcomes for consumer poultry products in this supply chain than those typically observed in the industrial food system, particularly with regard to the prevalence of multi-drug resistant (MDR) foodborne pathogens. This study had four specific aims: (1) describe the prevalence of *E. coli, Salmonella spp.,* and *S. aureus* in a market-basket sample of raw poultry meat purchased in the Maryland direct-market poultry supply chain; (2) characterize the antimicrobial resistance phenotypes of any isolates detected by culture; (3) compare these outcomes to relevant food safety data from National Antimicrobial Resistance Monitoring System (NARMS) and other independent peer-reviewed research; and (4) use matched data obtained with a survey tool from the same participating farms and poultry processors to explore associations between farm characteristics and observed food safety outcomes.

## 2 Methods and Materials

### Enrollment and Recruitment

We identified participants via publicly-available commercial registries that promote direct-market agricultural producers in Maryland, particularly the databases maintained by University of Maryland Agriculture Extension program [42]. As a secondary strategy, we used snowball sampling [18] to identify participants whose contact information was not available through the aforementioned sources. Participants were recruited via email or phone contact and offered a $20 cash incentive. The lead author conducted all of the surveys at the farms or homes of participants, and purchased a sample of frozen poultry at the conclusion of each survey. The Johns Hopkins Bloomberg School of Public Health Institutional Review Board approved this project and participants provided written informed consent for survey and oral consent for meat sampling.

### Survey tool

We administered a survey questionnaire to a broad sample of Maryland direct-market poultry producers. The questions in the survey tool focused on descriptive characteristics and workplace practices of small-scale poultry production and processing models. A copy of the survey is included in the supplement. These factors included: scale and size of production and processing operations; professional experience of producers and processors; antimicrobial usage in poultry production; maintaining multiple animal species in close or overlapping proximity; sanitary practices during slaughter and processing; poultry production practices; use of on-farm and third-party processing facilities; and sourcing of livestock. On-farm processing refers to slaughter and processing operations that are constructed on the farm where the broiler poultry are raised, and exclusively process the birds raised on that farm. Third-party processors refers to slaughter and processing operations that process broiler poultry for a fee for other poultry producers. Other information gathered using the survey questionnaire included county-level location data and processor certification status under Maryland Department of Agriculture (MDA) or the United States Department of Agriculture (USDA). Data from each survey questionnaire was matched to a unique poultry sample’s microbial outcome data. Information from the survey were used to create categories for comparing microbial outcomes among different groups of vendors.

### Sample collection, transport and storage

All 40 survey respondents provided oral consent to submit a single poultry meat sample from their retail store for microbial analysis, and were recruited into the market-basket stage of this research. Previous research indicated that frozen products were the most common products marketed by this population [17]; only frozen products were obtained for microbial assessment. Frozen poultry samples were transported by cooler and were not allowed to thaw during transport to the laboratory freezer, where samples were stored at −20°C to await microbial culture.

### Microbial culture and antimicrobial susceptibility testing methods: Salmonella spp

Laboratory culture methods for *Salmonella spp.* were adapted from NARMS protocols for culture-based methods for retail meat surveillance [19]. Packages of frozen meat were set out in open coolers in the lab 12-16 hours in advance and allowed to warm to room temperature. Thawed packages were opened aseptically using sterile surgical instruments, then two 25 gram aliquots of surface muscle tissue, skin, and fat were removed aseptically, weighed and placed into a stomacher bag containing either 200 ml of double-strength lactose broth (Becton Dickinson-Difco) or 200 ml of 0.9% saline solution. Both aliquots were agitated and vigorously shaken for 60 seconds, then 15 ml of the rinsate from the aliquot in the lactose broth was pipetted into a sterile centrifuge tube, vortexed, and incubated overnight at 35°C. Fifty milliliters of rinsate was then pipetted from the aliquot in saline solution and vortexed with 50 ml of double-strength lactose broth in a sterile flask and the contents were mixed thoroughly. Fifteen milliliters of this mixture was pipetted into a sterile centrifuge tube and incubated for 24 hours at 35°C with the tubes from the enrichment broth stomacher bag. From each tube, 0.1 ml was pipetted into 9.9 ml of Rappaport-Vassiliadis medium (BD-Difco) and incubated for 16-20 hours at 42°C. One milliliter of these enrichment broths was transferred to 10 ml tubes of pre-warmed M-broth (BD-Difco) and incubated at 35°C for 6-8 hours. The broth mixtures were allowed to cool to room temperature and 10 μl were streaked onto Xylose Lysine Deoxycholate (XLD) agar plate (Becton-Dickinson) and incubated overnight at 35°C. After 24 hours, plates were examined for colonies typical for *Salmonella* growth (pink colonies with or without black centers). Any typical colony was streaked to a trypticase soy agar plate supplemented with 5% defibrinated sheep’s blood (Thermo Scientific-Remel) to confirm isolate purity. Culture-positive isolates were confirmed and tested for antimicrobial susceptibility using the BD Phoenix system. A list of the antimicrobials tested is included in the supplement.

### Microbial culture and antimicrobial susceptibility testing methods: E. coli

Laboratory culture methods for *E. coli* were adapted from standard food safety literature [19, 22, 23, 24, 25]. Packages were allowed to thaw and opened as described above, and a 25 gram aliquot of mixed tissue types was aseptically removed, weighed and placed in a sterile stomacher bag with 200 mL of MacConkey enrichment broth (MAC broth) (Becton-Dickinson) and shaken vigorously for 60 seconds. Fifteen milliliters of this rinsate was pipetted into a sterile centrifuge tube and incubated 16-20 hours at 35°C. Tubes were vortexed thoroughly, and 10 μl from each tube was streaked onto MacConkey agar (MAC agar) (Becton-Dickinson) plates, which were incubated 16-20 hours at 35°C. Where *E. coli*-like growth (round pink colonies with or without a dark center and a hazy area surrounding colonies) was observed, a single colony or a 1 μl loop of typical but overcrowded growth was streaked to a fresh MAC agar plate and incubated 16-20 hours at 35°C. Culture-positive isolates were confirmed using the BD Phoenix automated microbiology system for species identification and antimicrobial susceptibility testing [20, 21] at the Johns Hopkins Hospital Clinical Diagnostic Microbiology Laboratory.

### Microbial culture, antimicrobial susceptibility, and molecular testing methods: S. aureus

Laboratory culture methods for recovery of *S. aureus* isolates from poultry meat samples were adapted from food safety literature on recovery of poultry livestock-associated *S. aureus* and MDR-*S. aureus* [26, 27, 28]. Packages of meat were allowed to thaw and aseptically opened as described above. A 25 gram aliquot of mixed tissue was removed, weighed and placed in a stomacher bag with 200 ml of Mueller-Hinton Broth (Becton Dickinson) supplemented with 6.5% NaCl (MHB+). The bag was vigorously shaken for 60 seconds, then 15 ml was pipetted to a sterile centrifuge tube, vortexed, and incubated 16-20 hours at 37°C. Tubes were vortexed after incubation and a 10 pμl loop of enrichment broth was streaked to blood agar plates (Thermo Scientific-Remel) and incubated 24 hours at 37°C. Plates were examined for typical *S. aureus* colonies (shiny, round, grey/white and with or without hemolysis) and either a single colony (when present) or a 1 μl loop of typical growth was streaked to a Baird-Parker agar plate (Becton-Dickinson) and incubated 24 hours at 37°C. Plates were examined for typical growth of coagulase-positive staphylococci (round, grey/black colonies demonstrating lecithinase activity) and culture-positive samples were confirmed and tested for antimicrobial susceptibility using the BD Phoenix system. A list of antimicrobials tested is included in the supplement.

Molecular testing was preformed on presumptive staphylococcal isolates by PCR to confirm presence of the *S.* aureus-specific nuclease gene (*nuc*) [29]. Additional PCR assays were used to detect presence or absence the *mecA* or *mecC* genes encoding methicillin resistance [30], Panton-Valentine Leukocidin (PVL) genes *lukF-PV* and *lukS-PV* [31, 32], and the staphylococcal complement inhibitor (scn) gene [33]. Real-time quantitative fluorescence PCR assay (TaqMan PCR) was used to detect genes encoding staphylococcal enterotoxins A, B, C, and D (SEA, SEB, SEC, and SED) of *S. aureus.* [34]. Staphylococcal protein A *(spa)* typing was performed using the Ridom Staph Type standard protocol (http://www.spaserver.ridom.de/) and Eurofins Genomics sequencer (eurofinsgenomics.com).

### Laboratory Quality Control

Quality control was assessed for laboratory bias or error by use of positive and negative controls. Positive controls and laboratory blanks (uninoculated broth samples run through the culture protocol) each were deployed at a rate of 10% for the culture protocols of all three target species (4 blank samples and 4 ATCC-positive samples per species). ATCC 25922, ATCC 14028, and ATCC 25923 were used as positive controls for *E. coli, Salmonella spp.* and *S. aureus,* respectively.

### Statistical Analysis

We used matched survey data obtained from the same sample population, derived variables from these data and applied them to a regression analysis and a variety of nonparametric tests of association to predict outcomes for microbial contamination and antimicrobial resistance. Logistic regression and nonparametric tests were used to assess the strength and statistical significance of any relationships between the variables derived from the survey data and the binary outcomes associated with different measures of microbial contamination status. Simple and multiple logistic regression tests, along with non-parametric analyses were used to assess inter-group differences between different categories of poultry vendors, as well as the effects of freezing time on recovery of target microbes.

### Comparison to the National Antimicrobial Resistance Monitoring System

The National Antimicrobial Resistance Monitoring System (NARMS) is a federal surveillance system that has been in existence since 1997 to detect antimicrobial resistant bacteria that contaminate retail meat in the United States [35]. In this analysis, the NARMS dataset was utilized as an external comparison group for comparison to bacteria isolated in this study. As such, prevalence was analyzed with most comparable group: *E.coli* isolates cultured from retail poultry meat purchased within Maryland in the year 2014. [35].

## 3. Results

### Enrollment and Recruitment

Between October, 2014 and March, 2015 we identified and attempted to contact 93 potentially-eligible participants. Sixteen potentially eligible participants identified using this system did not respond to two separate messages left on business phone voicemails. Sixteen other respondents informed us that their operation was currently out of business and 11 respondents reported that they were no longer marketing poultry meat as part of their business. From the remaining 50 eligible participants, four declined to participate in the study, citing privacy concerns, and six more participants were unable to schedule a time to participate during the recruitment window. Ultimately, a sample of 40 eligible poultry farmers in Maryland identified through our recruitment process participated in the study. This process is outlined in Figure 1.

**Figure 1:**
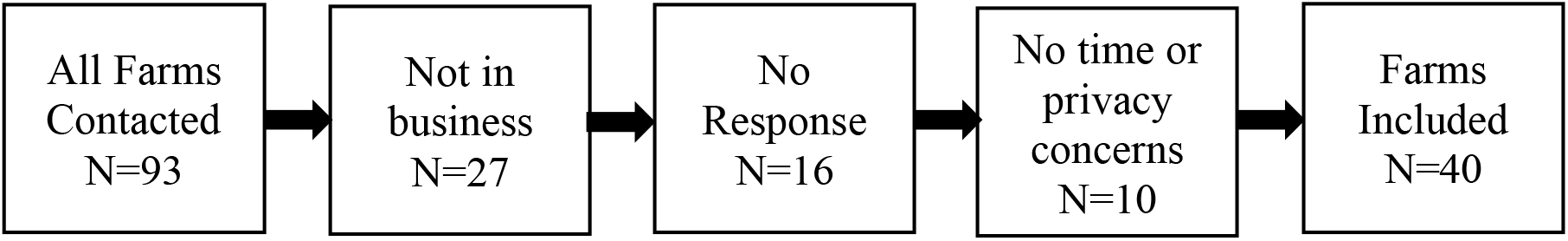
Flowchart for Enrollment and Recruitment of Participants

### Demographics and background information

Responses to the survey questionnaire were recorded and analyzed. Demographic information collected indicated that a majority (60%) of participants were female and 100% were white/Caucasian. Participants reported a median value of 5.5 years of professional experience, with an interquartile range of 2.5-10.0 years of experience. Figure 2 shows the geographic distribution of participating poultry farms at the county level across the state. Table 1 contains information on the scale of poultry production and on-farm practices among survey respondents, with most respondents indicating that they practiced on-farm poultry processing with a median flock size of 1,050 birds per year. Figures 3-6 summarize survey responses on the number and variety of other livestock and companion animals living on the same property as the poultry flocks. The vast majority of poultry production among respondents occurs in settings where poultry interact with and share a living environment with other livestock and companion animal species. Table 2 describes the sanitation and disinfection practices employed by respondents using on-farm poultry processing systems, indicating that a large majority of participants use two or more methods of disinfection both before and after a run of poultry slaughter and processing.

**Figure 2:**
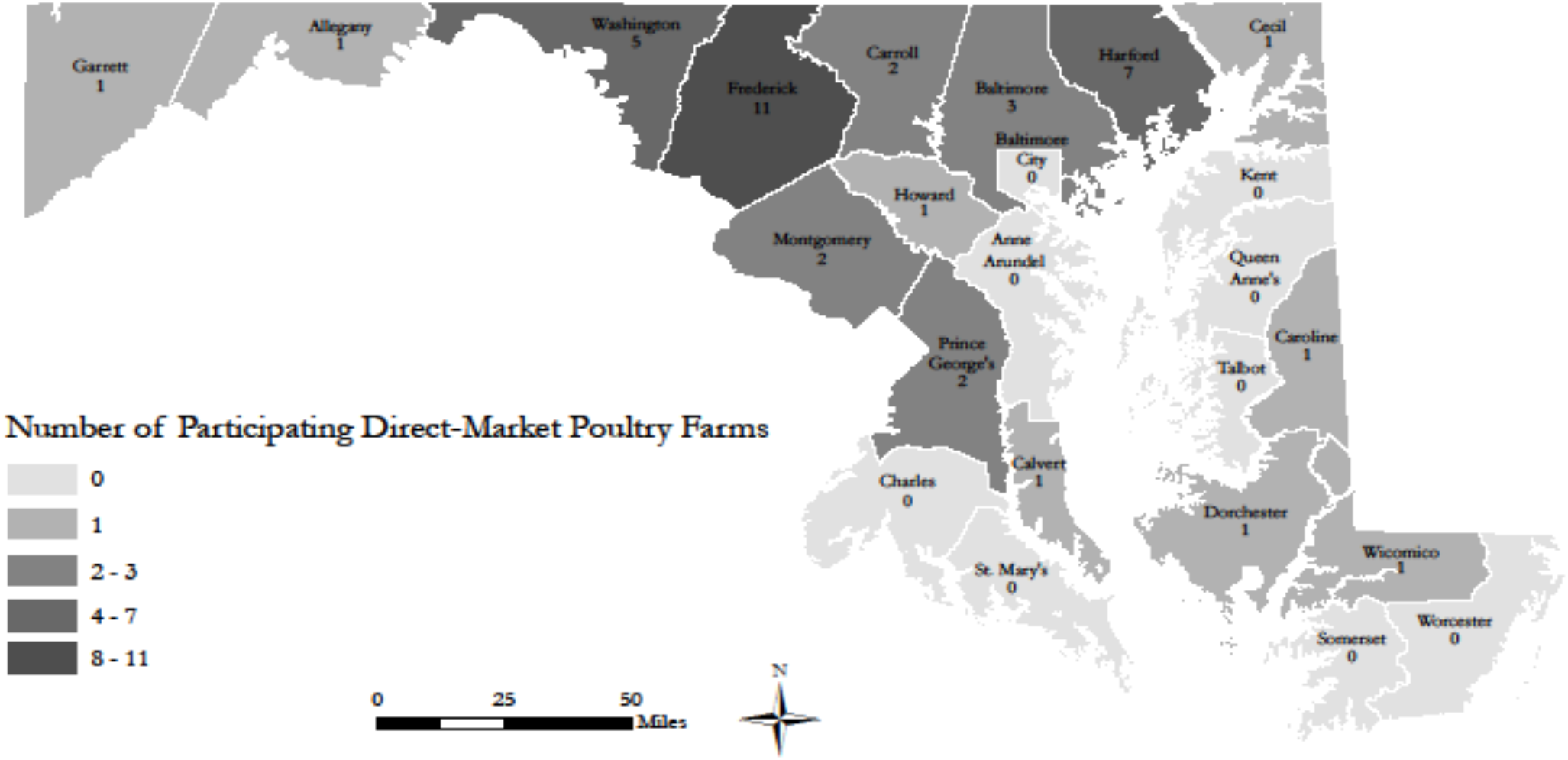
Geographic Distribution of Participating Poultry Producers in Maryland Counties

**Figure 3:**
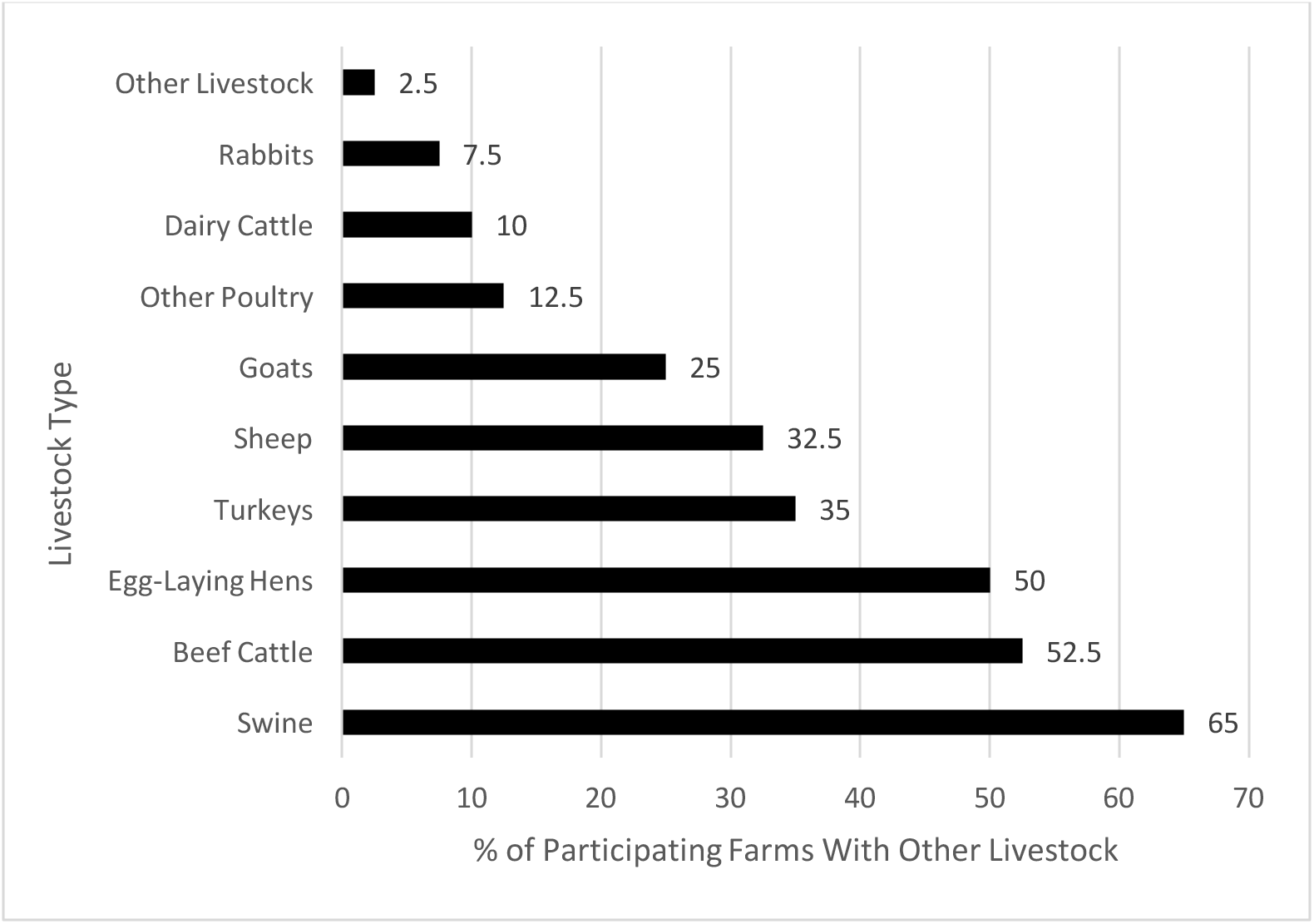
Percent of Participating Broiler Poultry Farms Keeping Other Livestock on Premises, By Type of Livestock

**Figure 4:**
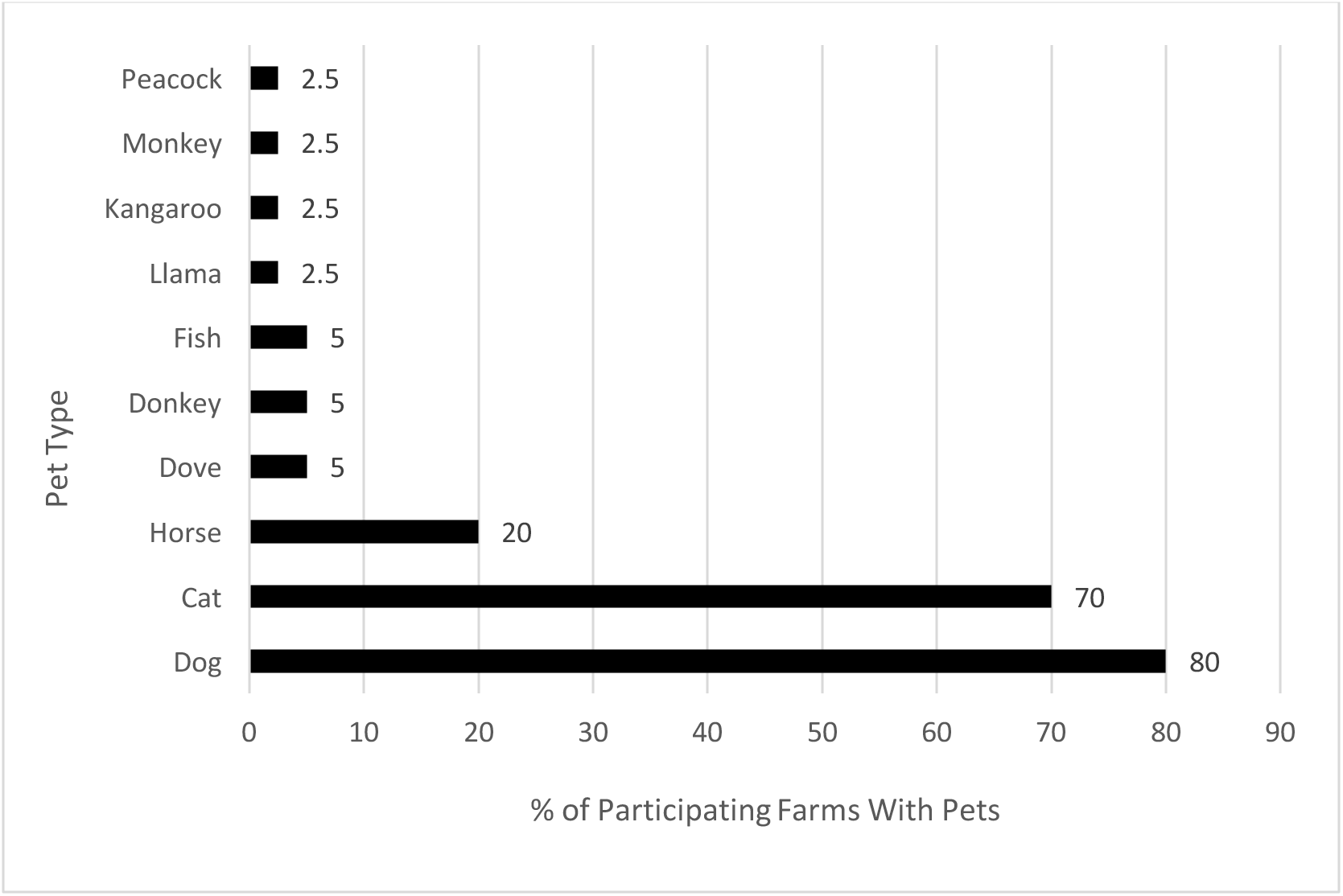
Percent of Participating Broiler Poultry Farms Keeping Pets on Premises, By Type of Pet

**Figure 5:**
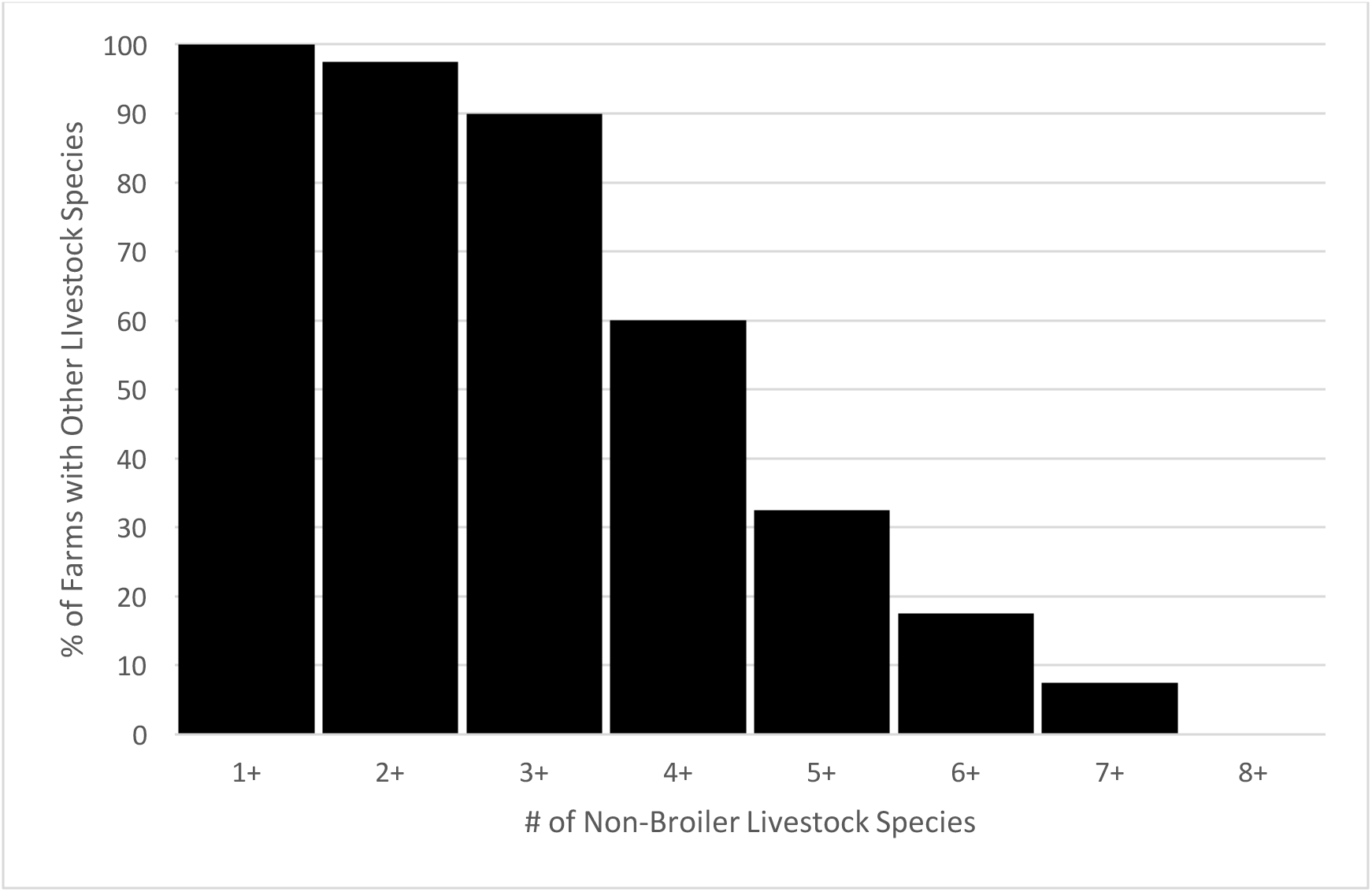
Percent of Participating Farms With Non-Broiler Poultry Livestock Species on Premises, by Number of Other Species

**Figure 6:**
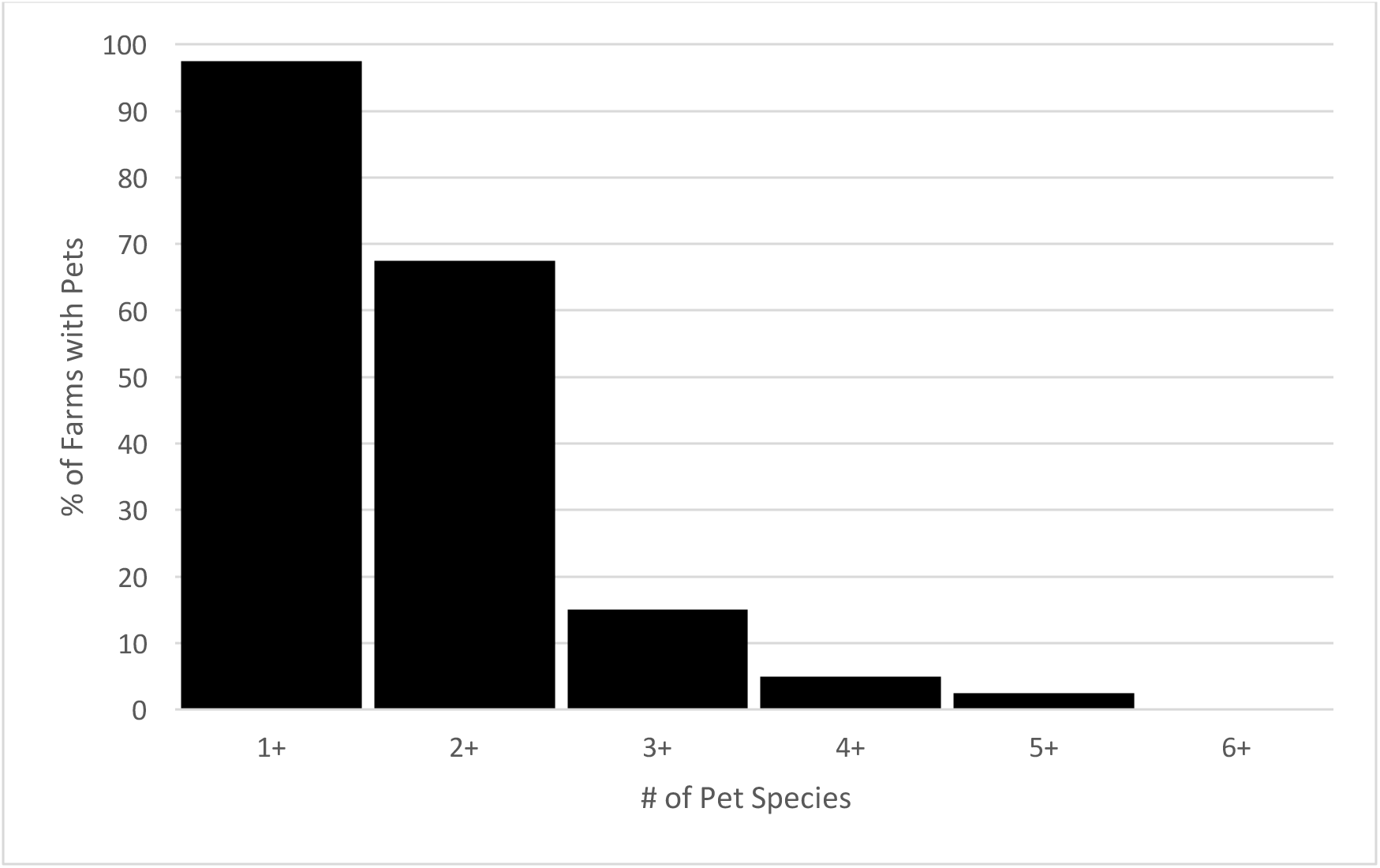
Percent of Participating Farms With Pets on Premises, by Number of Pet Species

**Table 1:** Production Scale of Maryland Direct-Market Poultry Operations

| Flock size | Total<br>(N=40) | On-Farm Processing<br>(N=25) |  |  | Third-Party Processing<br>(N=15) |  |  |
| --- | --- | --- | --- | --- | --- | --- | --- |
|  |  | USDA<sup>a</sup><br>(N=1) | MDA<sup>b</sup><br>(N=18) | Neither<br>(N=6) | USDA<br>(N=11) | MDA<br>(N=3) | Neither<br>(N=1) |
| Median (IQR)<sup>c</sup><br>Birds/yr | 1,050<br>(450-1,700) | 2,700<br>(2,700) | 1,200<br>(800-2,500) | 800<br>(200-1,700) | 800<br>(400-1,200) | 250<br>(150-800) | 2,000<br>(2,000) |
| Median (IQR)<sup>c</sup><br>Flock Size | 100<br>(50-150) | 200<br>(200-200) | 70<br>(50-100) | 33.5<br>(30-40) | 150<br>(100-175) | 100<br>(50-150) | 500<br>(500) |

a: USDA: United States Department of Agriculture Food Safety Inspection Service Certified Food Animal Processor b: MDA: Maryland Department of Agriculture Food Safety Certified Food Animal Processor c: IQR: Inter-Quartile Range

**Table 2:** Disinfection Practices of Participating OFPP Facilities, By Processing Stage (N=25)

|  | Pre-Processing (%) | During Processing (%) | Post-Processing (%) |
| --- | --- | --- | --- |
| Any Disinfection | 100% | 44% | 100% |
| Bleach Solution | 84% | 24% | 76% |
| Soap Water | 72% | 16% | 48% |
| Hot Water | 12% | 4% | 16% |
| Vinegar/Peroxide | 12% | 4% | 8% |
| UV (Sunlight) | 4% | 4% | 12% |
| No Soap, No Bleach | 4% | 76% | 16% |
| 2+ Cleaning Agents | 76% | 16% | 64% |

A minority (17.5%) of respondents reported using pharmaceutical antimicrobial inputs in poultry production. Among the 7/40 participants who used these inputs, three reported using antibiotics only to therapeutically treat sick livestock; all three reported exclusively using tetracycline administered through drinking water. The remaining four participants all reported preventative usage limited to recently-arrived chicks, who receive feed supplemented with coccidiostat drugs, and are put onto non-medicated feed for the “grow-out” period of production (from between 2-3 weeks to when the birds reach slaughter weight at ~7-12 weeks of age). Coccidiostats were the only antimicrobial inputs for which respondents reported prophylactic use.

### Prevalence and antimicrobial susceptibility of Gram-negative target species (E. coli, Salmonella spp.)

*E. coli* was recovered from 9/40 (22.5%) of retail poultry samples. Among the nine confirmed isolates, two were resistant to one class of antimicrobials; one isolate was resistant to tetracycline and the other to imipenem, a beta-lactam/carbapenem antibiotic. No *E. coli* isolates were resistant to more than one class of antimicrobials. Prevalence and antimicrobial-resistance phenotypes of *E. coli* among retail meat samples purchased from different categories of direct-market vendors is included in Table 3. Results comparing prevalence rates of AMR phenotypes among *E. coli* isolates recovered from 2014 NARMS surveillance in Maryland to the market-basket samples in this study are displayed in Table 4. No positive *Salmonella* isolates were recovered from any of the retail poultry samples analyzed in this study. The dual culture protocols that used either a lactose enrichment broth or 0.9% saline media as an initial aliquot did not yield differential results.

**Table 3:** Prevalence of AMR *S. aureus* and *E. coli* Isolates by Processor Location (on-farm vs. third party facility) and Food Safety Agency Inspection Status

| Processor Type | *S. aureus* | *S. aureus*:<br>1+ Classes<br>AMR | *S. aureus*:<br>2 Classes<br>AMR | *E. coli* | *E. coli*:<br>1 Class AMR |
| --- | --- | --- | --- | --- | --- |
| All Processors<br>(n=40) | 30% | 15% | 2.5% | 22.5% | 5% |
| 3rd Party<sup>a</sup> (n=15) | 33.3% | 13.3% | 6.7% | 26.7% | 6.7% |
| OFPP: All<sup>b</sup><br>(n=25) | 28% | 16% | 0% | 20% | 4% |
| OFPP: MDA/USDA<sup>c</sup><br>(n=19) | 21.1% | 15.8% | 0% | 26.3% | 5.3% |
| OFPP: Uncertified<sup>d</sup> (n=6) | 33.3% | 16.7% | 0% | 0% | 0% |

<sup>a</sup>: Vendors using a third party poultry processor;
<sup>b</sup>: All on-farm poultry processors (OFPP);
<sup>c</sup>: OFPP certified and by USDA(N=1) or MDA(N=18);
<sup>d</sup>: On-Farm Poultry Processor not certified by either/any agency;

**Table 4:** Prevalence of AMR in *E. coli* Isolates by Processor Location and Inspection Status from Sample Data Compared to 2014 Market-Basket Data from NARMS Surveillance in Maryland

| **Processor Type** | ***E. coli:***<br>**1+ Classes**<br>**AMR** | ***E. coli:***<br>**2+ Classes**<br>**AMR** |
| --- | --- | --- |
| All Processors<br>(n=40) | 5% | 0% |
| 3rd Party<sup>a</sup> (n=15) | 6.7% | 0% |
| OFPP: All<sup>b</sup><br>(n=25) | 4% | 0% |
| OFPP: MDA/USDA<sup>c</sup><br>(n=19) | 5.3% | 0% |
| OFPP: Uncertified<sup>d</sup> (n=6) | 0% | 0% |
| NARMS: Conventional<sup>e</sup><br>(n=166) | 77.1% | 63.9% |

<sup>a</sup>: Vendors using a third party poultry processor;
<sup>b</sup>: All on-farm poultry processors (OFPP);
<sup>c</sup>: OFPP certified and by USDA(N=1) or MDA(N=18);
<sup>d</sup>: On-Farm Poultry Processor not certified by either/any agency;
<sup>e</sup>: NARMS 2014 surveillance data of conventionally produced (non-USDA Organic) Maryland broiler poultry. Bacterial susceptibility was defined using minimum inhibitory breakpoints for *E.coli* with Clinical and Laboratory Standards Institute, M100: Performance Standards for Antimicrobial Susceptibility Testing, 29<sup>th</sup> Edition.

### Microbial prevalence and antimicrobial susceptibility of Gram-positive target species (S. aureus)

*S. aureus* was recovered from 12/40 (30%) of poultry samples. Of the 12 positive isolates, 6/12 were resistant to one or more antimicrobial classes, 1/12 were resistant to two antimicrobial classes, and none were resistant to three or more antimicrobials. All AMR *S. aureus* were exclusively resistant to tetracycline, penicillin and/or ampicillin. No multi-drug resistant *S. aureus* or methicillin-resistant *S. aureus* were recovered, and no *mecA* or *mecC* genes were detected. Four isolates were positive for the *scn* gene, which is a potential marker of human (rather than animal) origin. The *pvl* gene was not detected in any samples. No staphylococcal enterotoxin (SE) genes were detected in any samples. Eight unique spa-types where identified across the 16 isolates tested. The AMR phenotypes of all *S. aureus* recovered from poultry samples are displayed along different categories of direct-market vendors in Table 3 and in the heat map in Table 5.

**Table 5:**
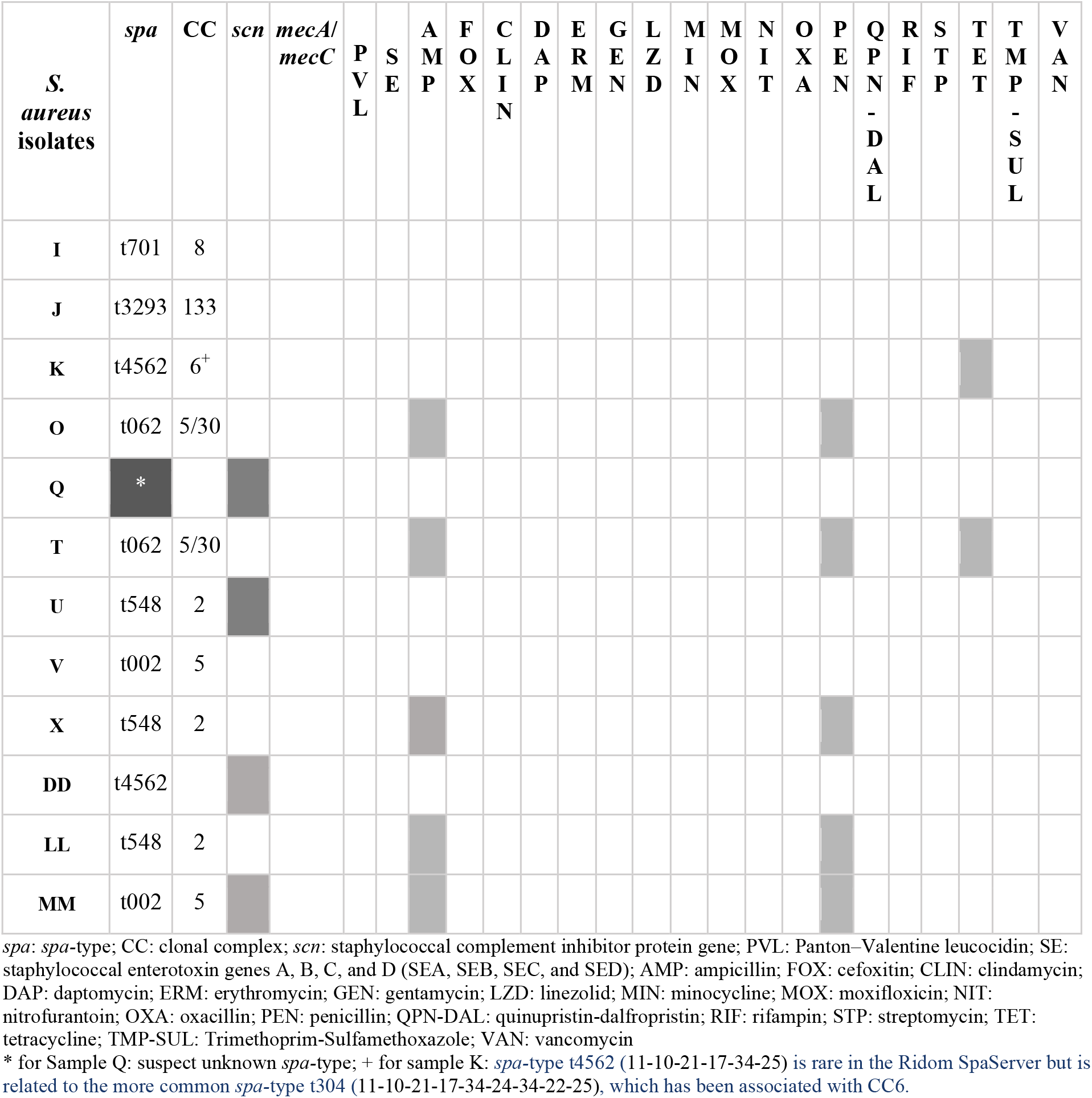
*spa*-types and Antimicrobial Resistance Among S. aureus positive isolates (n=12)

### Sample freezing time and regression analysis

Data used to calculate the duration of time between when the poultry carcass was processed and frozen and when the samples was thawed for analysis was available for 30/40 samples. For the remaining 10 samples this information was not on the label and could not be estimated accurately by the vendor. The samples had been frozen for an average of 140 days, with a range of 54-260 days and an interquartile range (IQR) of 108-150 days. Freezing time was treated as a continuous predictor variable for a simple logistic regression analysis for the outcome of finding any contamination, was used to determine a trend-level (*p*=0.08) increase in the odds ratio of finding any contamination with a one-day increase in freezing time (1.02, 95% CI: 0.99-1.04). This value was lower (1.01, 95% CI: 0.99-1.02) and the association was weaker (p=0.14) when the microbial outcome was limited to *S. aureus*-positive samples. When 10-day increases in freezing time were used to create an ordinal predictor variable for recovery of any target microorganisms, there were only slight changes to the observed association (1.04, 95% CI: 0.94-2.09) and the association was not statistically significant at α=0.05 (p=0.09). When 30-day increases in freezing time was used as an ordinal predictor variable for the same outcome, a stronger signal (1.86, 95% CI: 0.82-4.17) was observed, but this association was not statistically significant at α=0.05 (p=0.09).

## 4 Discussion

Overall recovery rates of *E. coli* were low and no *Salmonella spp.* were recovered. The 30% prevalence of *S. aureus* was comparable with the observed prevalence in the industrial-scale poultry supply chain [36]. Rates of antimicrobial usage were low (17.5%) among producers in this study, which may explain the very low rates of AMR from the market-basket sample and lack of detection of multidrug resistance among recovered isolates. Elimination of antimicrobial inputs in poultry production has been shown previously to be associated with lower rates of contamination of retail meat products with MDR microbial pathogens [37].

The distribution of *spa*/CC type of the *S. aureus* isolates recovered in our market-basket sample was similar to the distribution of isolates recovered from industrial market-basket samples of poultry and other meat products. Thapaliya *et al.* demonstrated t002/CC5 as the most prevalent *spa*/CC type among *S.aureus* isolates from their market-basket sample, recovering this type from ~15% of retail meat samples purchased in grocery stores in Iowa, USA. Approximately 17% of the *S. aureus* isolates from our market-basket sample were identified as t002/CC5; however, t548/CC2 was the most frequent *spa/CC* type identified, accounting for 25% of *S. aureus* isolates from our study sample.

### Survey Results

The survey data presented here quantify the frequency and characterize the distribution of structural elements and workplace practices of direct-market poultry operations that had been previously identified by research carried out in this population as important or relevant to microbial food safety [17]. Antimicrobial input usage was very low among participants; what usage was reported occurred under different conditions than those understood to drive the propagation of MDR foodborne pathogens in the industrial poultry supply chain. Only 10% of respondents from the direct-market supply chain reported use of antimicrobial inputs for disease prophylaxis in poultry flocks. Moreover, the antimicrobial inputs used by these respondents included only a single coccidiostat. Further, the antimicrobial mechanism associated with this drug is understood to be only weakly (if at all) associated with acquired AMR in bacterial populations [38]. None of the observed AMR phenotypes in our sample occurred in the samples from survey respondents reporting use of antimicrobial inputs for disease prophylaxis in their poultry flocks.

### Prevalence and AMR of target pathogens

The absence of MDR *E. coli* or *S. aureus* is a finding of particular public health significance. These results are strong supporting evidence for the hypothesis that some of the characteristics of direct-market poultry production may correlate with much lower prevalence of detection of drug-resistant *E. coli* on consumer poultry meat products (5%) compared to products from industrial poultry production (77.1%), based on NARMS surveillance data limited to poultry meat purchased in Maryland in 2014. *S. aureus* is not assessed routinely via NARMS surveillance [39].

The observed prevalence of *S. aureus* (32.5%) in this market-basket sample of Maryland direct-market retail poultry is roughly equivalent to trends observed in the few market-basket studies assessing the industrial poultry supply chain. This indicates that *S. aureus* is likely to still be a relevant food safety concern for direct-market poultry production. However, the absence of MDR *S. aureus* presents a major potential difference in the overall food safety health risks associated with this supply chain.

The absence of *Salmonella* positive isolates among the market-basket samples is surprising. Our negative results do not necessarily indicate an absence of viable *Salmonella* on these samples or within this supply chain. We can identify three possibilities that may explain these findings: (1) *Salmonella* concentrations were below the LOD of our methods; (2) freezing poultry reduced the viable number of *Salmonella*; (3) viable *Salmonella* isolates were present, but were injured or metabolically damaged by freezing and did not grow on selective culture media.

The rates of *E. coli* contamination are substantially lower than those reported in NARMS and in other research literature. In 2015, 63.5% of retail poultry meat samples sampled under NARMS surveillance were positive for *E. coli* contamination, similar to recovery of *E. coli* the prior year [39]. This may indicate a difference in food safety risks for consumers of direct-market products to be infected with fecal-origin bacterial contaminants, but more research is needed to establish the validity of those findings. As with *Salmonella,* freezing may play a role in reduction of *E. coli* recovered using these methods. Research on this topic within the industrial poultry supply chain has been limited and inconclusive as to whether different methods of freezing result in significant reductions in viable and recoverable *Salmonella spp.* and *E.coli* [40, 41].

### Strengths, limitations and areas for further research

One strength is of the study is having a mixed-methods approach that included both microbial sampling and survey interviews with participants. A second strength is that, while the study population was small, it captured ~60% of the population of direct-market poultry producers in Maryland and therefore these findings likely are generalizable to the entire population of Maryland producers.

There are several limitations, one being the sample size (N=40), which is small for a multiple logistic regression analysis. A second limitation was the cross-sectional study design—repeated samples would improve our ability to assess prevalence of microbial pathogens in the statewide direct market supply chain. Further, this study did not conduct serovar analysis of *E. coli* isolates or collect data to determine pathogenicity. In contrast, *S. aureus* isolates were tested for several characteristics related to pathogenicity, including presence of common enterotoxin genes linked to foodborne intoxication. In particular, sampling only frozen poultry samples presents both strengths and limitations to our analysis. Frozen poultry is the product form that consumers would purchase; however, freezing may affect target pathogen recovery. Fresh poultry products constitute the majority of samples in market-basket studies of the industrial poultry supply chain, which limits our ability to compare directly with these studies. Future research on this topic should address these limitations and seek to differentiate between pathogenic and non-pathogenic *E. coli* contamination of market-basket products, and consider to include additional poultry-associated foodborne indicator bacteria and pathogens, such as *Enterococcus* and *Campylobacter*.

This research is an important step to characterize the microbial food safety of food products from direct market poultry, which is an alternative to conventional poultry supply chains sold in supermarkets. These data provide evidence to support the potential for management practices that limit antimicrobial inputs to be associated with lower recovery of drug-resistant indicator bacteria and pathogens. These findings provide a baseline for future research on direct-to-consumer poultry products in Maryland and beyond, and may inform larger efforts to describe the contribution of food animal production to the global burden of drug-resistant pathogens.

## Acknowledgements

This research was carried out with laboratory support from Dr. Karen C. Carroll, MD and the Medical Microbiology laboratory at Johns Hopkins Hospital for microbial isolate analysis using the BD Phoenix automated microbiology system for species identification and antimicrobial susceptibility testing. Funding from the Johns Hopkins Center for a Livable Future’s Fellowship program supported this study. The funders had no role in study design, data collection and interpretation, or the decision to submit the work for publication. Dr. Christopher Heaney served as primary advisor to this project.

