## Supplemental Materials for "Microbial Food Safety in the Maryland Direct-to-Consumer Poultry Supply Chain"

The following is a copy of the survey tool administered to study participants (N=40)

Long Form Questionnaire

**Study Information**

This is a questionnaire designed to gather data on small poultry processors and retailers engaged in direct-marketing of poultry products in Maryland. The purpose of this study is to learn more about the range of models and practices that are currently being used to produce and process poultry in the state. We hope to apply some of this information to better understand what factors contribute to food safety in small poultry processing.

My name is____________________. I am a researcher with Johns Hopkins and the Center for a Livable Future, and I am conducting this survey only for research purposes. No personal information about you or your business will be collected as a part of this study. To contact us with any inquiries or concerns please email or call:

Patrick Baron

(410)-916-2112

**Recruitment question**

Hello, my name is ______________, I am a researcher from Johns Hopkins and the Center for a Livable Future, and I am gathering data about the practices used in raising and slaughtering the poultry you are selling. I would like to purchase some of your meat and to pay you an additional $10.00 cash for about 15 or 20 minutes of your time to complete a short survey about your poultry operation. Your responses will be completely anonymous. Would you be interested in scheduling a time today, or anytime soon that might work for you, to participate in a brief survey about the practices your farm uses to raise and slaughter poultry?

**Consent question**

Would you be interested in participating in this survey?

If **YES**, proceed to **Oral Consent Form**, if **NO**, proceed to **Refusal Questionnaire**

Once oral consent is obtained, the investigator must sign and date below and the survey may proceed.

Investigators Name (Print)________________________________

Investigators Signature____________________________________

Date_______________

Matched with Meat Sample:____________________

*The following questions are about your farm and your poultry processing business. They should take one to two minutes to complete.*

**Farm Characteristics:** Background and Inclusion/Exclusion criteria

1) Do you raise broiler poultry for commercial slaughter and sale?  (y/n)

- If **YES**, for how many years? ____________

2) Do you process poultry for commercial sale? (y/n)

    - if **YES**, for how many years?______

3) Do you raise other livestock animals for commercial slaughter and sale? (y/n)

- if **YES**, which types of animals and could you estimate how many of each type per year?

________________________________________________________________________________________

________________________________________________________________________________________

________________________________________________________________________________________

________________________________________________________________________________________

4) Do you keep pets or other non-livestock animals on your farm that are not raised for commercial slaughter? (y/n)

- if **YES**, which types of animals and could you estimate how many of each type are typically present on your farm?

________________________________________________________________________________________

________________________________________________________________________________________

________________________________________________________________________________________

________________________________________________________________________________________

*The next questions are about your poultry processing operation as well as some of the characteristics of how the poultry you sell was raised before slaughter. They should take about five to ten minutes to complete.*

**Poultry Processing General Questions**

5. Was the poultry you have for sale today processed on your farm or at a facility owned and operated by you personally? (y/n)

If **YES,** proceed to Questions 6-18, if **NO**, proceed to Question 19

6. Is your poultry processing facility certified under the MDA’s Rabbit and Poultry Slaughter and Processing Training Program for FSIS-Exempt Producers? (y/n)
7. Is your poultry processing facility certified by the USDA? (y/n)
8. Where the facility that processes your poultry located? (Check all that apply)
   a. Completely outdoors
   b. Partially outdoors (shed or temporary structure)
   c. Indoors (in a building only accessible from the outside by doors and windows that can be completely closed)
9. Could you estimate how many birds you typically process during a single day of slaughtering? _________________________(#)
10. Do you ever slaughter broiler poultry **from other producers** at your facility? (y/n)

If **YES,** proceed to Question 11, if not, skip to Question 12

11. Could you estimate how many birds **from other producers** you slaughtered in the past 12 months? ________________________ (#)
12. Could you estimate how many of **your** birds you slaughtered in the past 12 months? _________________________ (#)
13. Do you typically slaughter poultry on consecutive days? (y/n)
14. Do you process other animals or game besides **broiler** poultry at your facility (check all that apply)
    - Turkeys
    - Ducks
    - Geese
    - Other poultry____________
    - Rabbits
    - Deer
    - Other game______________
    - Other animals_____________

*The next questions will be about the equipment and workplace methods you use for processing poultry at your facility. Some questions will be directed towards the practices you use to clean your facility and processing equipment and to keep poultry carcasses clean. This section should take about five minutes to complete.*

**Poultry Processing Equipment and Methods**

15. When you typically process poultry, do you clean **any** of the processing equipment and surfaces:
    a. before beginning the run? (y/n)

               If **YES:**

i. Which equipment or surfaces do you clean?* ________________________________________________________________________________________________________________________________________________________

____________________________________________________________________________

ii. Which disinfectants to you use to clean these?

________________________________________________________________________________________________________________________________________________________

____________________________________________________________________________

**many participants simply say “everything”, “contact surfaces” or “contact and non-contact surfaces” instead of individual equipment or surfaces. You can just record these responses as-is.*

b. during processing?* (y/n)

               If **YES:**

i. Which equipment or surfaces do you clean? ________________________________________________________________________________________________________________________________________________________

____________________________________________________________________________

ii. Which disinfectants to you use to clean?

________________________________________________________________________________________________________________________________________________________

____________________________________________________________________________

**NOT during a contamination event, just under normal, typical circumstances of processing*

c. after the run is completed? (y/n)

              If **YES:**

i. Which equipment or surfaces do you clean? ________________________________________________________________________________________________________________________________________________________

____________________________________________________________________________

ii. Which disinfectants to you use to clean?

________________________________________________________________________________________________________________________________________________________

____________________________________________________________________________

16. Do you use any disinfectants to clean the carcasses during processing when/if you observe carcass contamination? (y/n)

If **YES**, answer question 17, if **NO**, skip to question 18**:**

17. Which disinfectants* do you typically use to clean contaminated carcasses?

_______________________________________________________________________________________________________________________________________________________________________________________________________________________________________________________________________________________

**water rinse or hot water wash is a frequent response, record as-is*

18. When you typically process poultry, how many of the following people (including yourself) participate in the poultry processing run at your facility?
    a. full-time employees _______  (#)
    b. part-time employees _______ (#)
    c. volunteers or family members_______ (#)

**Questions for participants who use third-party processors**

*Skip this section (Questions 19-23) if the participant responded* **YES** *to Q5; e.g. if they own/operate the processing facility that processed the poultry we are sampling. Skip to Question 24.*

19. Was the poultry you have for sale today processed at a facility operated by a third party? (y/n)
20. Has the facility that processes your poultry been certified under the MDA’s Rabbit and Poultry Slaughter and Processing Training Program for FSIS-Exempt Producers? (y/n/don’t know)
21. Has the facility that processes your poultry been certified by the USDA? (y/n/don’t know)
22. Could you estimate how large the typical flock of broiler poultry that you bring to be slaughtered and processed at this facility is? ______________________________(#)
23. Could you estimate how many of your birds were processed at this facility in the last 12 months?_____________________________(#)

**Characteristics of the flock(s) you process**

24. How many hatcheries have supplied live chicks to your farm in the last year? (1,2,3,4,5,6,7,8,9,10+)
25. Are these hatcheries NPIP*-certified? (yes/no/don’t know)

*Circle all that apply if these answers apply to more than one hatchery.*

**Note: NPIP = USDA National Poultry Inspection Program*

26. Are the poultry slaughtered in your facility raised in permanent housing structures, or are mobile housing structures in use?*
    a. Permanent
    b. Mobile
    c. Both
    d. Do not know

**Many pastured poultry farmers use a permanent housing structure for baby chicks, and during the last 6-8 weeks before processing the birds are moved to mobile housing structures. If the farmer describes this scenario, mark “Both” as the response.*

27. If mobile chicken housing is used, how frequently is the location of the housing moved during production?
    a. Daily
    b. 2-3x week
    c. Weekly
    d. Bi-weekly
    e. Monthly
    f. Less frequently
    g. Do not know
    h. Mobile housing not in use
28. Are pharmaceutical antibiotics or antimicrobials used in poultry production for any of the birds that you slaughter for sale? (y/n/do not know)

If **YES**, answer Questions 29-31, if **NO**, questionnaire is complete**:**

29. Antibiotics are used in these manners: (Check all that apply)
    a. to promote growth/feed efficiency in the absence of disease
    b. to prevent disease (sometimes called prophylactically)
    c. to treat an infection or disease outbreak
30. Through what route are the antibiotics administered (check all that apply)
    a. In feed
    b. In water
    c. By injection
31. Which antibiotics or antimicrobial drugs do you use? (choose all that apply)
    a. Amoxicillin
    b. Cefovecin
    c. Cephalexin
    d. Clavulanic Acid
    e. Doxycycline
    f. Enrofloxacin
    g. Neomycin
    h. Moxifloxacin
    i. Marbofloxacin
    j. Ofloxacin
    k. Penecillin
    l. Pirlimicin
    m. Streptomycin
    n. Sulfasalazine
    o. Synulox
    p. Trimethoprim
    q. Tylosin
    r. Ionophores
    s. Coccidiostats
    t. Other______________________

*End of Survey*

Supplemental analysis of Scheinberg *et al.* 2013

This single study, which sampled whole fresh (n=60) and frozen (n=40) chicken carcasses purchased from 21 vendors at different farmers’ markets in Pennsylvania, evaluated the prevalence of non-typhoidal *Salmonella* and *Campylobacter* on whole carcasses, and compared these results to comparable microbial data obtained from analyzing USDA-Organic and conventional poultry meat at Pennsylvania supermarkets. The study demonstrated very high prevalence (85% and 93% for frozen and fresh samples, respectively) of *Campylobacter* contamination for poultry purchased from farmer’s market vendors, significantly higher than conventional (52%) or Organic (28%) chicken samples. Prevalence of non-typhoidal *Salmonella spp.* was reported to be statistically significantly higher in farmer’s market poultry than for conventional poultry products (28% and 8%, respectively), and higher than Organic poultry meat (20%). While this research is a first step in filling in the knowledge gap for food safety issues in direct market poultry supply chains, this study contains (and significantly, does not report) obvious limitations to interpretations of the study data. No power or sample size calculations are reported for assessing statistical significance of any of the reported associations; and there is no description of an attempt to assess a study population N (total number of direct-market poultry producers in Pennsylvania) to determine the representativeness of the pool of study participants (N=21 vendors). Relevant data that can generally be obtained from the label of the consumer product, such as whether the bird was slaughtered and processed at a USDA-FSIS inspected facility, a state-inspected facility, or an uninspected on-farm processing facility, is not reported. The study assumes that all vendors practice on-farm poultry slaughter and processing under FSIS-exemption, but does not verify or report this important information, which may not be accurate in many cases. Our survey of Maryland direct-market poultry vendors found that over 37% of these producers use a third-party processor that is not located on their farm, and that 30% processed their broiler poultry livestock at a USDA-FSIS inspected facility. Finally, this study does not assess antimicrobial-resistance characteristics of foodborne pathogens in their market-basket sample. Given the critical significance of antimicrobial resistance in zoonotic and foodborne pathogens for food safety and public health in general, this is a major omission in any attempt to characterize microbial food safety risk issues in the direct-market poultry supply chain.

BD Phoenix Panels

BD Phoenix System Antimicrobial Susceptibility Panel (Gram-Negative)

Amikacin, Amoxicillin/Clavulanate, Ampicillin, Ampicillin/Sulbactam, Aztreonam, Cefazolin, Cefepime, Cefotaxime, Cefoxitin, Ceftazidime, Ceftriaxone, Cefuroxime, Ciprofloxacin, Ertapenem, ESBL Confirmatory, Gentamycin, Imipenem, Levofloxicin, Meropenem, Moxifloxacin, Nalidixic Acid, Nitrofurantoin, Piperacillin, Piperacillin/Tazobactam, Tetracycline, Ticarcillin/Clavulanate, Tobramycin, Trimethoprim/Sulfamethoxazole

BD Phoenix System Antimicrobial Susceptibility Panel (Gram-Positive)

Ampicillin, Ampicillin/Sulbactam, Cefazolin, Cefoxitin, Cephalothin, Chloramphenicol, Clindamycin, Daptomycin, Erythromycin, Gatifloxicin, Gentamycin, Gentamycin-synergy, Levofloxicin, Linezolid, Moxifloxacin, Nitrofurantoin, Norfloxacin, Oxacillin, Penicillin, Quinupristin/Dalfopristin, Rifampin, Streptomycin, Tetracycline, Trimethoprim/Sulfamethoxazole, Vancomycin
